# Scopolamine induced learning deficit in marmosets

**DOI:** 10.1101/2025.11.18.688868

**Authors:** Haley E. Harkins, Karen Christopher, Denis Matrov, Ivan D. Ingram, Eric B. Saglio, George R. Dold, Yogita Chudasama

## Abstract

In monkeys, the muscarinic cholinergic receptor antagonist scopolamine is known to broadly disrupt learned behaviors, though the precise nature of the cognitive deficits has been questioned. Experimentally observable deficits in memory can be ascribed to poor attentional focusing, human interference as well as age, sex, and dosing regimen. Stress and social isolation can also play a role during behavioral testing, particularly in small nonhuman social primates like marmosets that have been used widely. In this study, we examine the effects of scopolamine in marmosets under conditions of reduced stress, attentional distraction, and human interference. Using a custom designed home-cage touchscreen-based testing system, we investigated the influence of scopolamine on the performance on a visual associative learning task. During self-paced, voluntary testing, monkeys learned to discriminate pairs of complex visual patterns through trial and error by touching the stimulus associated with reward. Using this approach, we demonstrated over 75% discrimination accuracy in the eight marmosets tested (male and female) within three days of home-cage testing. Although the averaged data revealed no impact of acute or chronic scopolamine injections on learning, modeling the choice data with trial-level analysis revealed both age- and sex-specific deficits. The results demonstrate the value of home-cage testing combined with trial-level analysis to reveal subtle behavioral changes, such as those brought about by scopolamine.

**Significance statement:** We created a custom home cage testing system to test the effects of muscarinic cholinergic blockade on complex discrimination learning in marmoset monkeys. We found that systemic scopolamine administration disrupts visual association learning in a manner that was specific to older females. This deficit was hidden in session-averaged measures and only became evident in trial-level modeling of the choice data. Our findings demonstrate that cholinergic blockade impairs the dynamics of learning in marmosets and highlights the value of trial-level analysis for detecting nuanced pharmacological effects on primate cognition.

## Introduction

Scopolamine is a muscarinic receptor antagonist which readily crosses the blood-brain barrier and inhibits the action of acetylcholine in the central nervous system. In humans, it has long been known to have deleterious effects on the formation of long-term memories and disrupts the process of memory consolidation (Drachman and Leavitt, 1974; Broks et al., 1988; Robbins et al., 1997; Edginton and Rusted, 2003; Green et al., 2005). Animal studies have also shown that scopolamine disrupts performance in a variety of learning and memory tasks (Beatty and Bierley, 1985; Aigner et al., 1991; Givens and Olton, 1995; Melamed et al., 2017). Much of this work derived from monkeys tested on automated discrete-trial delayed response tasks to understand memory-related deficits associated with Alzheimer’s disease (Bartus et al., 1978; Aigner and Mishkin, 1986; Spinelli et al., 2006). However, the precise nature of the performance deficits have been questioned, since scopolamine can also affect non-mnemonic functions (Drachman and Leavitt, 1974; Petersen, 1977; Beatty et al., 1986; Goldman Rakic, 1987; Pang et al., 1993; Muir et al., 1994; Voytko et al., 1994; McGaughy et al., 1996), including spatial perception, attention or response perseveration (Mishkin and Manning, 1978; Callahan et al., 1993; Chudasama and Muir, 1997). Moreover, scopolamine-induced cognitive deficits can be transient, dose-dependent, and can depend on age and sex (van Haaren et al., 1989; van Hest et al., 1990; Gallagher and Colombo, 1995). This variation in results has made it difficult to tailor cholinergic pharmacotherapies for psychiatric conditions such as schizophrenia or mood disorders whose symptoms are closely linked to dysregulated muscarinic receptors (Foster et al., 2021; Paul et al., 2022; Yohn et al., 2022).

In the present study, we used adult marmoset monkeys (*Callithrix jaccus*) to assess the effect of scopolamine on a visual associative learning task that does not require a memory load. Ridley and colleagues previously showed in marmosets that scopolamine does not generally impair memory, but instead affects the acquisition of newly learned information (Ridley et al., 1984a; Ridley et al., 1984b; Harder et al., 1998). In addition to its central effects, scopolamine can have adverse effects on parasympathetic activity including pupil dilation, tachycardia and dehydration, symptoms which can worsen in stressed animals. Since marmosets are a social species, laboratory practices of individual testing of animals causes social isolation and raises stress cortisol levels, and this may impact the normal expression of behavior (Norcross and Newman, 1999; Cross et al., 2004; Kaplan et al., 2012). Moreover, the release of acetylcholine in brain regions can vary considerably depending on environmental conditions and behavioral demands (Klinkenberg et al., 2011; Micheau and Marighetto, 2011; Mineur et al., 2022). Thus, the systemic administration of scopolamine might be expected to depend on the specific testing conditions, sometimes showing deficits (Kirk et al., 1988; Drinkenburg et al., 1995; Bushnell et al., 1997; Robinson et al., 2004), and sometimes not (Cheal, 1981; McGaughy et al., 1994; Harder et al., 1998). This combination of factors likely explains the inconsistent behavioral effects of scopolamine observed in various species (Fibiger, 1991; Terry Jr, 2006; Klinkenberg and Blokland, 2010).

In this study, we approached this problem with a home cage system allowing marmosets to perform self-paced, voluntary testing. Many laboratories have capitalized on home cage testing to reduce the impact of unknown confounding factors such as stress or human interference on behavior (Savage et al., 1987; Crofts et al., 1999; Takemoto et al., 2011; Calapai et al., 2022; Glavis-Bloom et al., 2022; Scott et al., 2025). Each day, the monkeys voluntarily entered the home cage testing box and proceeded to successfully discriminate several pairs of visual stimuli to a high degree of accuracy. After they had learned the task, we challenged the animals with acute and chronic doses of scopolamine and applied both session-average and trial-level analyses to assess the effects on visual discrimination learning.

## Materials and Methods

### Subjects

Eight adult common marmosets (*Callithrix jacchus*), 3 males and 5 females, ages ranging from 2-9 years served as subjects in the present study. Animals were exposed to a 12:12 light: dark cycle with artificial light illumination (300 LUX) and natural light from a window. Room temperature was maintained at 22-23 degrees Celsius and 65-68% humidity. Cages were changed every two weeks, and the enrichment structures inside the cage were changed as needed. All marmosets had free access to food and water except 1-3 hrs before testing. They were fed a daily diet of commercial marmoset food that was modified (Test Diet 5WW6) for high fiber and gum Arabic to support digestive health (Mazuri Marmoset Diet, St. Louis, MO), supplemented with fresh fruit and vegetables, nuts, mealworms, and caterpillars. All procedures accorded with the Guide for the Care and Use of Laboratory Animals and were approved by the NIMH Animal Care and Use Committee.

### Apparatus

All testing was conducted in four custom designed home-cage operant boxes (**Fig. 1A**). The boxes were adapted to connect to the animal’s home cage such that once securely mounted, a sliding door, when opened, allowed the monkey to enter the operant box. The sliding door remained open to allow the animal free access to the home cage during the entire test schedule. Each box was constructed from a combination of Plexiglass and steel and comprised two compartments. One compartment was integrated with the touchscreen and accessible to the animal (12″ x 13.25″ x 12″). Located opposite the touchscreen was a touch-sensitive licker which dispensed 0.2ml banana milkshake (∼3 drops) as a reward. Opaque grey blinders attached to the clear surface of the box occluded distraction from others monkeys in the homeroom. The second compartment (6″ x 13.25″ x 12″) housed the Arduino Uno microcontroller (R3, Code #A000066, DigiKey, USA), lithium-ion battery pack (Talentcell.com, NB7102), and touchscreen cables. The home-cage operant box was controlled by the Arduino microcontroller integrated with either a Dell Latitude 7210 or 7320 laptops running PsychoPy version 2022.2.5.

**Figure 1.**
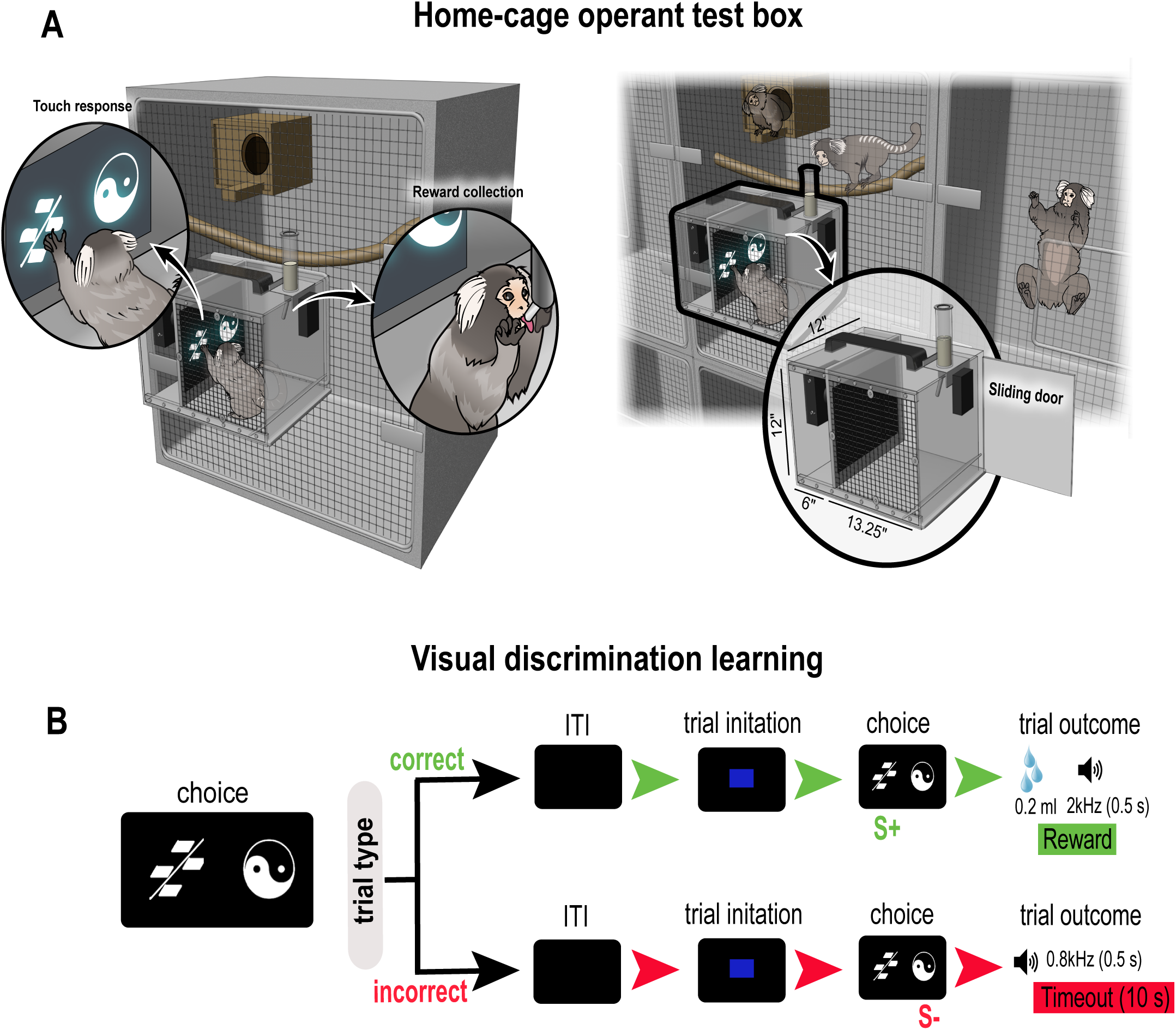
Home-cage touchscreen operant testing box and visual discrimination task. **A.** Schematic illustration of a home-cage operant test box attached to a home cage. Magnified images in left panel shows marmoset viewing/touching visual stimulus presented on a touchscreen and collecting reward from a licker tube. Magnified test box in the right panel shows location of sliding door which when opened, allows the monkey to enter and exit the box. **B.** Schematic illustration of visual discrimination task showing sequence of trial events for correct and incorrect trial types. After an intertrial interval (ITI), which varied between monkeys depending on how often they returned to their home cage, marmosets initiated each trial by touching a centrally positioned blue square which resulted in the presentation of the choice stimuli (correct: S+ or incorrect: S-). Touching the correct stimulus delivered 0.2ml reward concomitant with a 2kHz sound for 0.5sec. An incorrect choice resulted in a blank screen for 10 sec, a 0.8kHz for 0.5sec and no reward.

### Behavioral procedure

We tested animals on a visual discrimination task (**Fig. 1B**) to assess fundamental aspects of forming stimulus-reward associations (Schultz and Dickinson, 2000), and because this task has been commonly used with marmosets to understand the underlying cholinergic mechanisms important for cognitive function (Voytko et al., 1994; Ridley et al., 1999; Spinelli et al., 2006). Animals were first allowed to habituate to the home cage test box for 2 hrs a day and then shaped to associate a tone (2kHz) with reward delivery from the licker tube (∼2 weeks). They were then trained to touch the screen (touch training, **Fig. S1A**). Initially, the entire screen was touch sensitive so that any touch on the screen was rewarded. We first presented a blue rectangle which covered 100% of the screen (10″ x 12″). A touch anywhere on the blue screen triggered the licker to release 0.2 ml reward concomitant with a 2kHz tone. Over successive days, the size of the blue rectangle and the touch sensitive area were gradually reduced to 50% so it was centrally located on the screen (2.3″ x 2.3″) to hone the animal’s dexterity and touch response within the touch sensitive area. Touch training sessions were 2hrs long per day and took six days to complete **(Fig. S1B-D).** When animals were reliably touching the stimulus that occupied only 50% of the screen and completing approximately 50 trials, they were ready for discrimination learning.

In discrimination learning, each trial began with the presentation of a blue square (3.7″ x 3.7″) in the center of a black screen. A single touch to the blue square resulted in its disappearance and the presentation of two computer graphic stimuli on the left and right side of the screen (**Fig. 1B**). One stimulus was designated the correct S+ and the other, the incorrect S-, counterbalanced across animals. The left and right positions of the stimuli were determined pseudorandomly. A touch to the correct stimulus resulted in the disappearance of the stimuli, and the delivery of 0.2 ml reward concomitant with 2kHz tone for 0.5 seconds. A touch to the incorrect stimulus resulted in a 10 sec timeout with no reward and 0.8kHz tone for 0.5 sec **(Fig 1B**). The monkeys routinely returned to their homecage before initiating the next trial so the intertrial interval (ITI) was determined by each animal and their motivation to initiate the next trial.

Initially, we used two distinguishable stimuli (stimulus pair 1) to establish a learning criterion. This revealed that over a 2 hr session period, monkeys readily completed an average of 100 trials achieving on average 75% accuracy within 3 days (**Fig. 2**). This stimulus pair served as the baseline data for the acute scopolamine injections. Subsequently, new stimulus pairs were used depending on the drug protocol described below.

**Figure 2.**
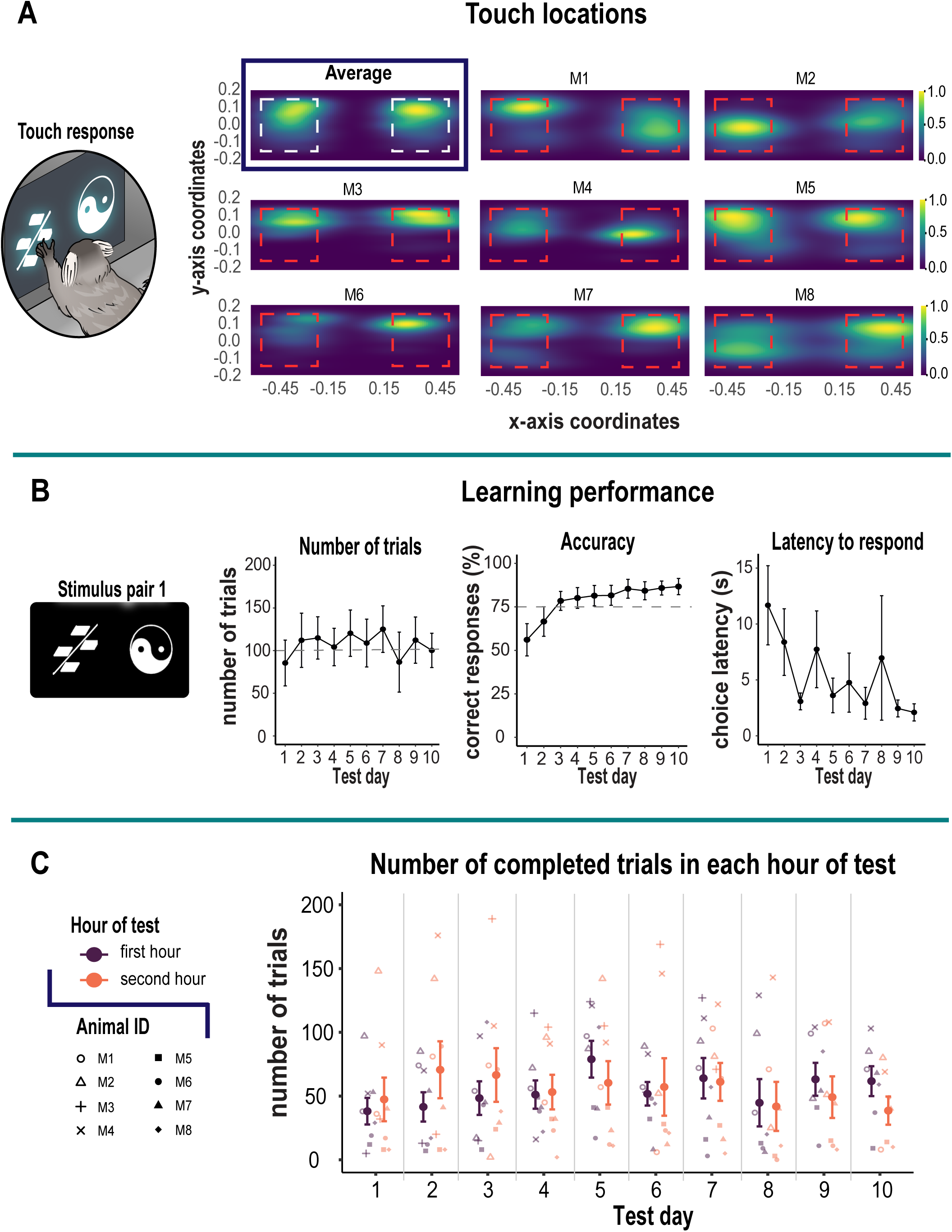
Acquisition of the visual discrimination task. **A.** Magnified image in left panel shows marmoset viewing/touching visual stimulus pair 1 presented on a touchscreen. Right panel shows combined 2D kernel density estimation of touch locations for all marmosets as an average, as well as the touch locations for individual marmosets which show horizontally elongated patterns. The red dashed square represents touch sensitive zone. Coordinates for touch locations were obtained from touch sensitive zone only. **B.** Left panel shows visual stimulus pair used for acquisition of discrimination task. Left graph shows number of trials (± S.E.M.) committed over 10 test days with an average criteria of 100 trials (gray dashed line). Center graph shows mean accuracy (± S.E.M.) across test days. The dashed gray line represents the 75% accuracy criterion. Marmosets attained 75% criteria by test day three. Right graph shows mean (± S.E.M.) latency to respond to the choice stimuli which decreased from 12 sec on test day 1 to 2 sec on test day 10. **C.** Graph shows the stability of the animals’ performance over the 2-hr session duration by quantifying the number of trials each animal completed in the first hour relative to the second as an index of motivation. Mean (± S.E.M.) is represented by the bold colors. Individual animals are represented by different shapes.

### Acute scopolamine injections

We administered single (acute) injections of scopolamine to examine its impact on the already learnt stimulus pair (**Fig. 3)**. Animals were first exposed to an intramuscular (i.m.) mock injection procedure for two days. On the following days, they received saline (vehicle), a low dose (0.03 mg/kg) or high dose (0.07 mg/kg) injection of scopolamine. Injections were given 1 hour before testing. Both vehicle and drug days were separated by a drug free day of no testing. Scopolamine hydrobromide (Sigma Aldrich, Pharmacopeia Reference Standard, Catalogue # 1610001, United States) was dissolved in sterile saline and prepared fresh before testing.

**Figure 3.**
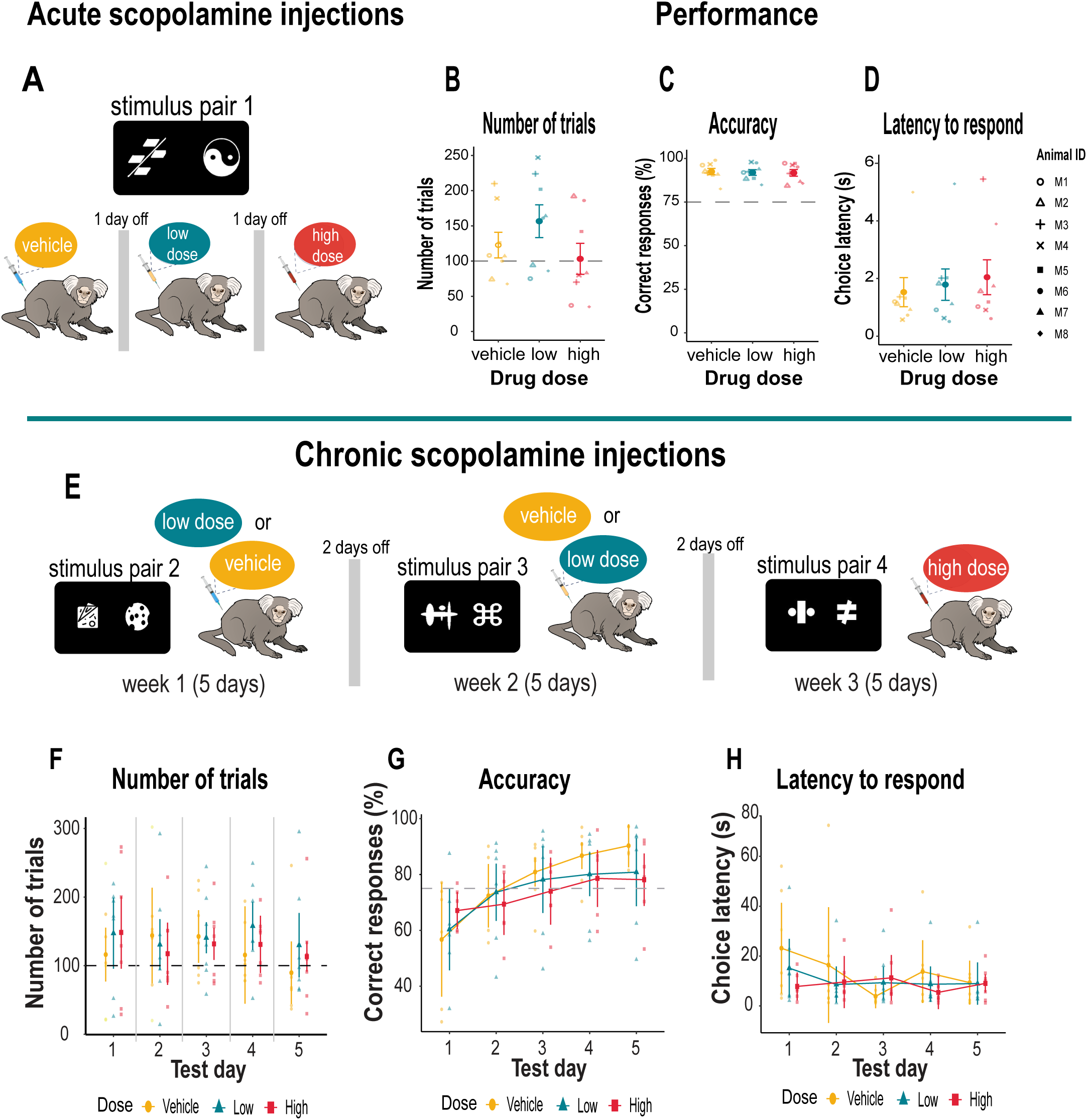
Experimental design and discrimination performance with scopolamine. **A.** Schematic illustration of experimental design for acute (single) injections of systemic scopolamine when discriminating stimulus pair 1. Animals received vehicle (saline), a low dose (0.03 mg/kg) and high dose (0.07 mg/kg) scopolamine on three separate days. **B.** Mean number of trials (± S.E.M.) committed under each drug condition with the average criteria of 100 trials (dashed gray line). **C.** Mean accuracy (± S.E.M.) on each drug injection days. Criterion was set at 75% accuracy (dashed gray line). **D.** Mean (± S.E.M.) latency to respond on each drug injection day. Individual animals are represented by different shapes. **E.** Top panel provides a schematic illustration of the experimental design for chronic (daily) scopolamine injections. Animals received vehicle (saline) or a low dose (0.03 mg/kg) scopolamine injection in a counterbalanced manner for five consecutive days (weekdays) over a two week period. The high dose (0.05 mg/kg) was lowered to prevent potential sedative side effects and was administered daily to all animals for five consecutive days. A different stimulus pair (pair 2, 3, 4) was presented each week. Graphs show 5-day acquisition of different stimulus pair when challenged with vehicle of scopolamine. Individual data points represent animals in each drug group. **F.** Mean number of trials (± S.E.M.) for each drug condition with the average criteria of 100 trials (dashed gray line). **G.** Mean accuracy (± S.E.M.) for each drug injection day. Criterion was set at 75% accuracy (dashed gray line). **H.** Mean (± S.E.M.) latency to respond on each drug injection day.

### Chronic scopolamine injections

We next introduced a chronic regimen of scopolamine. This time, animals were injected i.m. with vehicle, a low dose (0.03 mg/kg), or high dose (0.05 mg/kg) for five consecutive days (Monday to Friday) (**Fig. 3**). The high dose was lowered to avoid potential sedative effects of the chronic scopolamine regimen. A different stimulus pair was used each week for each dose so we could examine the effects of scopolamine on new learning. The drug was injected one hour before behavioral testing. For the first two weeks, the injections for vehicle and low dose scopolamine were counterbalanced so that half of the animals received vehicle on week one and low dose scopolamine on week two, and *vice versa*. For the high dose condition, all animals received the same dose for the whole week. Each drug condition was separated by two days. Each session lasted 2 hours.

### Data analysis: learning and acute scopolamine injections

The different performance measures for learning the visual discrimination and for the acute injections of scopolamine were subjected to a repeated measures ANOVA using IBM SPSS (version 30). Homogeneity of variance was assessed with Mauchly’s sphericity test with a Greenhouse–Geisser correction to provide a more conservative p value. The within-subjects factors was number of sessions (10 sessions for learning) and dose of drug at three levels for the acute scopolamine injections (vehicle, 0.03 mg/kg, 0.07 mg/kg).

### Data analysis: chronic scopolamine injection

The data were analyzed both in aggregate on a per-session basis, and on a per-trial basis to study predictors of correct task performance (i.e., logistic regression). We first filtered out trials where the number of touches outside the stimulus touch-sensitive zone exceeded 20. Long sequences of touches to the non-response area of the touchscreen indicated to us that the animal’s behavior was unfocused, impulsive and thereby unrelated to the discrimination. There were 16 such trials in total (M1 had eight such trials, M2 four, M8 three, and M7 one). We applied a repeated measures ANOVA with a within-subjects factors of session (five consecutive days) and dose of drug at three levels (vehicle, 0.03 mg/kg, 0.05 mg/kg). The statistical analysis was conducted in RStudio (version 2024.04.1) using the R statistical programming environment (version 4.3.3). A paired t-test was conducted using the *rstatix* package. The ANOVA was performed with *afex* package (version 1.3.1), and the statistical modeling of single trial data was done with *gamlss* package (version 5.4.22).

The single trial analysis allowed us to make use of the additional statistical power afforded by having over 15 thousand records of the individual trials and to fit the generalized linear models. We looked for conditional effects of a number of continuous covariates such as age or response latency, and factors such as sex. We used the Generalized Additive Model for Location, Scale and Shape (GAMLSS) (Stasinopoulos et al., 2017) framework to model the probability of making the correct discriminative choice by using the binomial distribution. The GAMLSS model assumes conditionally independent observations of the response variable given the model parameters, the explanatory variables, and subject-specific random effects. The data were unbalanced, as the number of completed daily trials varied between and within marmosets. To account for the different number of observations in each test day, as well as the serial correlation between subsequent observations, each trial datum was nested within the marmoset and test day variables. Such model specification treats the marmosets as random effects with flexible time-dependent trajectory. Moreover, to further account for serial correlation in the data, the sequences of testing days and trials were modeled using P-splines and treated as nuisance parameters. The predictors were checked for collinearity and the final model was chosen by minimization of the Akaike information criterion’s score and visual inspection of model diagnostic plots.

### Code Availability

All the code used for the home cage’s microcontroller (Arduino), the cognitive tasks from Psychopy, and the code used for statistical analysis and data processing is available for view and use at: https://github.com/SBNResearcher

## Results

We created a custom home-cage operant box integrated with a touchscreen that securely latched onto the animals home-cage (**Fig. 1A**). This afforded us the opportunity to assess marmoset cognitive abilities in an unrestrained environment within its home-cage vicinity. We found this approach maximized performance while avoiding the stress associated with daily capture. Animals successfully touched the stimuli located on the left and right of the screen. **Fig. 2A** shows the aggregated 2D kernel density estimation plot of touch locations for a learned stimulus pair that combines data from all marmosets. The plot shows no side bias. However, individual density plots for each animal revealed horizontally elongated patterns reflecting a strategy of swiping the stimulus in a horizontal direction using their hand.

In the context of the self-paced home cage testing apparatus, animals learned the stimulus reward discrimination within three days. On average, they completed 106 trials per day (range 85 – 125) within a 2-hr period as they learned the stimulus-reward contingency (**Fig. 2B, left panel)**. The criterion for learning was 75% accuracy, and by session 3, marmosets attained an average of 84%. The animals were then tested for an additional 7 days to ensure the behavior was stable over time. We found the animals maintained above-criterion levels of performance, which further improved with increasing number of test days [F_(9,63)_ = 5.82, p < 0.001]. By day 10, animals were attaining over 87% accuracy **(Fig. 2B, middle panel).** Consistent with their accuracy, the latency to make a choice became increasingly fast over the 10 day period [F_(9,63)_ = 3.65, p < 0.001). This measure was more variable over the course of testing because the animal had free access to its home-cage; often times, the monkey would return to its home-cage before completing the trial. Regardless, the speed of responding decreased from 12 to 2 sec by day10 **(Fig. 2B, right panel)**. We also examined the stability of the animals’ performance over the 2-hr session duration by quantifying the number of trials they completed in the first hour relative to the second as an index of motivation. On average, the animals performed 55 trials in each hour confirming their high motivation to work for the entire 2-hr duration (**Fig. 2C)**.

### Scopolamine does not affect previously learned stimulus associations

Early studies in monkeys report inconsistent findings concerning the impact of acetylcholine blockade with acute doses of scopolamine on learned behavior. While some suggest it disrupts the encoding of information into memory, thereby causing subsequent learning impairments (Evans, 1975; Ridley et al., 1984a; Harder et al., 1998), others report no such effect (Ridley et al., 1984b; Spinelli et al., 2006). Most of these studies were conducted manually using junk or random objects as stimuli, and monkeys were tested individually in specially designed operant boxes. We therefore examined if a single low (0.03 mg/kg) or high (0.07 mg/kg) dose of scopolamine would impact accuracy of a learned discrimination (stimulus pair 1) in our automated home-cage testing set-up which was void of experimenter interference and allowed the animal close proximity to its family **(Fig. 3A).** Under these conditions, neither dose of scopolamine had any impact on behavior (**Figs. 3B-D**). Specifically, the number of completed trials for vehicle and each drug dose was equivalent [F_(2,14)_ = 2.07, p = 0.16] as was performance accuracy [F_(2,14)_ = 0.14, p = 0.87] and their latency to make a choice [F_(2,14)_ = 1.12, p = 0.35]. This demonstrates that a single injection of a low or high dose of scopolamine does not impair post-acquisition learning or memory of the stimulus association, their perception or encoding of the stimuli, or their motivation.

### Scopolamine does not affect the learning of new stimulus associations

To rule out the possibility that monkeys overlearned the discrimination since the same stimulus pair was used before and after the scopolamine injections, we next tested the effects of more chronic administration of scopolamine on marmosets’ ability to acquire new stimulus discriminations. To limit the potential detrimental effect of scopolamine on the autonomic nervous system during the chronic dose regimen, the high dose of scopolamine was reduced to 0.05 mg/kg. Marmosets received injections of vehicle, low dose 0.03 mg/kg or high dose 0.05 mg/kg for five consecutive days. The vehicle and low dose regimen were counterbalanced over a two-week period (see **Fig. 3E)**. Each week, the animals were required to learn the stimulus-reward contingencies of a new stimulus pair. Although there was high variability across subjects, the dose of drug [F_(1.92, 11.53)_ = 1.12, p = 0.87] or test day [F_(2.68, 16.08)_ = 0.76, p = 0.52] did not affect the number of completed trials (**Fig. 3F**) with all animals completing an average of 135 trials a day. However, animal M1 repeatedly selected stimuli on the same side of the touchscreen indicating a side bias. This monkey was therefore excluded from the analysis of discriminative accuracy. The remaining subjects (n=7) showed no effect of scopolamine on side preferences (left side touches: p = 0.30, right side touches: p = 0.18). Moreover, the average number of consecutive touches for each stimulus presented on the left and right sides of the touchscreen were equivalent (p = 0.18).

Overall, the dose of scopolamine had no main effect on performance accuracy (F_(1.68, 8.38)_ = 0.28, p = 0.728), which improved across test days (F_(1.57, 7.87)_ = 17.91, p = 0.002), but a dose x test day interaction [F_(8, 40)_ = 2.95, p = 0.011] suggested that the impact of the dose was not consistent across days (**Fig 3G**). While the vehicle injections led to a steeper improvement in learning, performance accuracy was generally lower following scopolamine injections on test days 4 and 5 (**Fig. 3G**), but not statistically significant (p > 0.05). Response latency was also in the normal range for all doses across days [F_(1.94, 11.64)_ = 1.02, p = 0.39] and test day [F_(2.12, 12.72)_ = 1.33, p = 0.30] (**Fig. 3H**). We note, however, that although scopolamine had no major impact on the animals’ behavior, there was much inter-subject variability within our group thereby creating a heterogenous sample of data. We therefore applied trial-level analysis to examine the trial-by-trial variability within each subject to understand how individual differences might relate to the behavioral outcomes.

### Accurate performance with low dose scopolamine depends on sex and age

We investigated whether two important factors, sex and age, were important variables in the effects of scopolamine on task performance. The marmosets used for this study were an opportunistic sample comprising four young adults (2 years old) and four older ones (6-10 years), 3 males and 5 females. Due to limited statistical power, the effects of age and sex on discrimination accuracy could not be fully explored within the ANOVA framework. As an alternative analysis approach, we modeled the performance of individual animals based on single-trial data (see Methods). This was facilitated by the thousands of trials we collected over the course of testing. We used a standard method (Olejnik and Algina, 2003; Stasinopoulos et al., 2017) to model the probability of making a correct response with logistic regression. The same model was applied to two datasets: one included all trials from all animals and one with animal M1 removed since this animal showed a response bias described in the previous section. This let us check whether the results depended on that outlier animal. In both cases, the models provided a good fit to the data. **Fig. 4A** shows diagnostic plots for the model with the full dataset. These plots are based on residuals (the difference between observed and predicted responses). The far-left panel shows the residuals versus predicted probabilities to make a correct choice. Each point represents the residual for a single trial. Residuals are scattered around zero across the full range of predicted probabilities (0 to 1), with no clear pattern or trend. This indicates that the model predictions were not systematically biased at any probability level. The second panel shows how the residuals behave across trials. Each residual appears evenly spread around zero across all trials, without any clear trend or systematic drift over time. Thus, the residuals are fairly random and stable across trials. The third panel shows the distribution of the residuals as a smooth bell shaped curve, centered around zero. This supports the assumption of normality. The fourth panel shows another confirmation of a normal distribution, this time as a Quantile-Quantile (Q-Q) plot. Together, these plots confirm that the model assumptions were well met, and the model was a good fit to the data.

**Figure 4.**
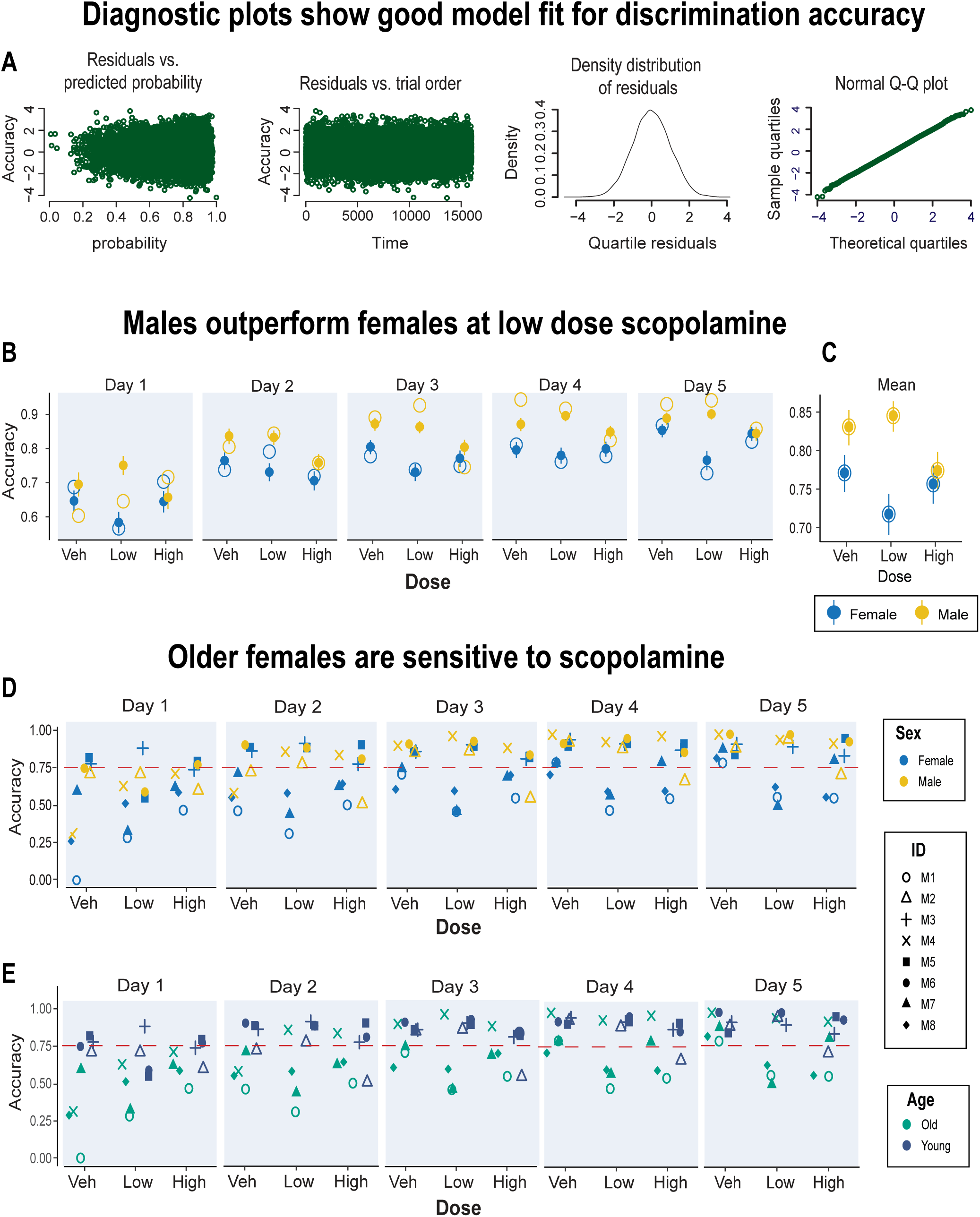
Trial-level analysis: scopolamine-induced performance depends on sex and age. **A.** Four diagnostic plots show that the model was a good fit for discrimination accuracy data (see text for details). **B.** The model-predicted correct choice probability stratified by the test day, dose, and sex, plotted as solid circles. The model predictions are presented as means with 95% confidence interval. The experimental data are plotted as hollow circles. **C.** Graph shows these data marginalized over the test days for each drug condition. **D.** Accuracy scores for each marmoset stratified by the drug dose and test day. The data points are mapped to color according to marmosets’ sex (**D**) and marmosets’ age (**E**). Marmosets above 6 years were classified as older monkeys. The dashed horizontal line corresponds to 75% criterion performance.

**Table 1** lists the model coefficients. The baseline group was female marmosets in the vehicle condition. The data are presented as odds ratios where coefficients greater than one indicate better odds of a correct choice, while values between zero and one indicate worse odds. For example, male marmosets had odds ratios greater than 1 relative to females, showing better overall performance. In contrast, chronic scopolamine treatment had odds ratios less than 1, indicating worse performance on discrimination trials. In addition to the main effect of sex and scopolamine, the interaction between these factors was also significant, indicating that the effects of scopolamine were both dose and sex dependent.

**Table 1.** Analysis of single trial data showing model predicted odds ratio coefficients of regressors on the probability of choosing the correct stimulus for datasets with animal M1 (full dataset) and without M1 who showed a response bias. The coefficients between 0 and 1 (exclusive) signify a negative relationship between the variable and the probability of choosing the correct stimulus. Conversely, the odds ratio of one corresponds to the absence of the relationship, and the coefficients larger than one postulate a positive association.

| Full dataset (15989 observations) |  |  |  | Dataset without M1 (14922 observations) |  |  |
| --- | --- | --- | --- | --- | --- | --- |
| Predictor | Odds Ratio | 95% CI | p-value | Odds Ratio | 95% CI | p-value |
| sex [male] | 1.26 | 1.07 – 1.49 | **0.006** | 1.15 | 0.97 – 1.36 | 0.099 |
| dose [low] | 0.65 | 0.57 – 0.73 | **<0.001** | 0.64 | 0.56 – 0.73 | **<0.001** |
| dose [high] | 0.83 | 0.73 – 0.94 | **0.004** | 0.81 | 0.70 – 0.93 | **0.004** |
| age | 0.88 | 0.87 – 0.90 | **<0.001** | 0.90 | 0.89 – 0.91 | **<0.001** |
| choice latency | 0.87 | 0.83 – 0.91 | **<0.001** | 0.90 | 0.85 – 0.95 | **<0.001** |
| correct response on previous trial | 1.00 | 0.91 – 1.10 | 0.985 | 1.13 | 1.03 – 1.25 | **0.013** |
| number of touches | 1.02 | 0.99 – 1.05 | 0.231 | 1.01 | 0.97 – 1.04 | 0.670 |
| distance to the stimulus' center | 1.72 | 1.65 – 1.80 | **<0.001** | 1.66 | 1.59 – 1.73 | **<0.001** |
| sex [male] × dose [low] | 1.72 | 1.39 – 2.13 | **<0.001** | 1.73 | 1.39 – 2.16 | **<0.001** |
| sex [male] × dose [high] | 0.77 | 0.62 – 0.95 | **0.014** | 0.80 | 0.64 – 1.00 | **0.045** |

One important revelation from the trial level analysis was that the effect of the low dose of copolamine on test performance was sex dependent; females performance was disrupted by scopolamine which, if anything, appeared to enhance the performance of the males (**Figs. 4B, C**). Thus, in general, males outperformed the females especially with low dose scopolamine. The individual data for discriminatory accuracy are plotted in **Fig. 4D**. This shows that this impairment was observed in a subset of animals. With vehicle, all monkeys attained over 75% accuracy by day 5. Even animal M1, who had a side bias, reached 79% accuracy. In contrast, the low dose of scopolamine reduced the accuracy of three marmosets to below 75%. These three individuals (M1, M7, and M8) were all females aged six years or older suggesting that older females were more sensitive to scopolamine (**Fig. 4E**). The small sample makes it difficult to further disentangle the independent effects of age and sex on these variables.

In addition, the age of animal and latency to make a choice were both negatively associated with performance indicating that older animals tended to perform worse, and slower responses were more likely to be incorrect. Interestingly, while the number of touches to the stimuli were not impacted in anyway, the location of the touch was relevant. Specifically, a positive association was observed between the Euclidean distance of the touch from the stimulus center and the probability of a correct choice suggesting that animals had a preference for a specific region of the stimuli when making their response.

## Discussion

We created a custom touchscreen operant apparatus to present complex visual discrimination tasks to marmosets in their home cages. All monkeys successfully performed a series of single pair visual discriminations with over 75% accuracy after 3 days of testing. The animals maintained a sustained level of interaction with the touchscreen actively participating in hundreds of trials, showing high levels of performance and became increasingly fast in their decision speed, which is normally recognized as an index of learning. Compared with traditional laboratory testing of marmosets in remote testing chambers (Ridley et al., 1981; Roberts et al., 1988; Clarke et al., 2015), the home cage operant system eliminated the stress associated with social isolation, the risk of handling the animal, experimenter interference, and made the procedure less labor intensive. These advantages, we believe, had a major influence on the marmosets rapid learning rate and consistent performance over time; they were highly motivated and worked reliably for each hour of the 2-hr session duration. The home cage testing system had the added advantage of providing cognitive enrichment; by allowing the animals to choose when to engage, it enabled self-directed discrimination learning, and they did it with high accuracy (Calapai et al., 2022; Calapai et al., 2023). The administration of scopolamine, however, had hardly any impact on learning ability; there was no systematic effect of scopolamine on the daily number of performed trials, their speed of response, or their accuracy in discriminating the stimuli. Moreover, there was no evidence of sedation, even at the high dose since performance effects of both high and low doses of scopolamine were very similar.

The repeated exposure of the *same* stimulus pair might have caused the animals to ‘overlearn’ the discrimination problem suggesting that the training continued beyond the initial point of learning. This might explain the high rate of discriminative accuracy (∼90%) following acute scopolamine injections. In humans and marmosets, scopolamine does not interfere with material learnt prior to drug treatment (Mewaldt and Ghoneim, 1979; Ridley et al., 1984a) such that once the learnt material is well encoded and retained in long-term storage, the animals performance is highly resistant to change. In the current study, however, scopolamine did not prevent the animals from acquiring ‘new’ discrimination problems even when it was administered daily for five consecutive days. Here, the animals rapidly transferred the learning rule to new discrimination problems even though the stimuli were different for each dose. This superior rate of discrimination learning suggests that the animals had acquired a *learning set* (Harlow, 1949; Miles and Meyer, 1956; Yokoyama et al., 2004). In other words, the animals had developed strategies and rules which they readily applied or transferred to the new stimulus pairs. Moreover, since all the new stimulus pairs were exemplars of the same dimension (i.e., shapes) it is likely that repeatedly being rewarded for choosing the same dimension led to an attentional bias or expectation that this dimension will continue to be relevant across new exemplars(Slamecka, 1968; Dias et al., 1996; Collins et al., 1998). This focused level of attention leads to faster learning and fewer errors. Previously reported scopolamine induced learning impairments in monkeys were specific to three dimensional objects that were perceptually complex (Evans, 1975; Ridley et al., 1984b; Harder et al., 1998). The visual stimuli in the present study were also perceptually difficult to discriminate because they were equated for luminance thereby preventing the marmosets from using brightness as a salient discriminating cue. We can be sure therefore, that monkeys were discriminating the stimulus features, but even this visual complexity was not enough to increase processing load since scopolamine failed to block the acquisition of learning new stimulus pairs. It is possible that the degree of muscarinic cholinergic receptor occupancy was insufficient to produce an effect (Yamamoto et al., 2011), but this seems unlikely, especially for the chronic dose conditions where the doses were comparably similar or higher to other studies (Ridley et al., 1984b; Spinelli et al., 2006; Melamed et al., 2017). Thus, from the aggregate data alone, there was no suggestion that learning under scopolamine was detrimental in any way to the animals’ behavior.

The advantages associated with the home-cage testing chamber might have facilitated the animals’ cognitive proficiency as shown by others (e.g., (Takemoto et al., 2015; Calapai et al., 2022), but we noted much variability between monkeys in their performance; like humans, these animals differed in their temperament, motivation and attention to the task. Compounded by the small sample size and differences in age and gender, we surmised that individual variability may have masked any meaningful change in behavior. At the same time, we had accrued thousands of trials per animal over many sessions. This allowed us to apply single trial analysis to capture trial-by-trial fluctuations in the animals’ performance and reveal subtle scopolamine-induced biological patterns of behavior. One main finding from this analysis was the sex-specific pattern in learning performance that was not apparent when the data were averaged across sessions. Generally speaking, female marmosets were more sensitive to the effects of scopolamine and therefore displayed lower gains in accuracy than males. In addition, the low performing females tended to be older females (> 6 years). Sex specific cognitive differences with scopolamine have been reported in animals with older animals exhibiting greater vulnerability (Ray et al., 1992; Tariot et al., 1996; De Castro and Girard, 2021). It is thought this reflects scopolamine’s interaction with estrogen which declines with increasing age (Voytko, 2002; Dumas et al., 2006; Dumas et al., 2010). Estrogen enhances cholinergic signaling, particularly in the basal forebrain and the hippocampus (Luine, 1985; Gibbs et al., 1994; Gibbs, 2010). In several rodent studies, estrogen can improve or protect against scopolamine-induced cognitive impairments, especially in ovariectomized rats (Fader et al., 1998; Gibbs et al., 2004; Tanabe et al., 2004). Since scopolamine is a muscarinic receptor antagonist, it may strongly counteract estrogen-dependent modulation in females. It is feasible therefore, that in the current study, scopolamine exacerbated estrogen-related cholinergic decline in older females which impaired their learning dynamics relative to males. Although our sample size reduces predictive power, our data are in keeping with several studies which report cholinergic associated cognitive decline in aged females typically exacerbated by loss of ovarian estrogens in both monkeys and humans (Hogervorst et al., 2000; Lacreuse et al., 2002; Rapp et al., 2003; Sherwin, 2006).

We cannot be sure of the exact locus of scopolamine action in our study, but studies that have directly targeted the medial septal/vertical diagonal band cholinergic projections to the hippocampus with excitotoxic or immunotoxic lesions confirm impaired visuospatial learning in adult marmosets (Ridley et al., 1988; Ridley et al., 1989; Harder et al., 1998). In contrast, lesions of the cortically projecting cholinergic neurons of the basal forebrain in rats and monkeys affect attentional processes (Roberts et al., 1992; Voytko et al., 1994; McGaughy et al., 2002; Chudasama et al., 2004; Dalley et al., 2004). Thus, presumably, the net effect of systemic dosing of scopolamine in the present study antagonized postsynaptic muscarinic receptors in the hippocampus. Consistent with this view, local hippocampal infusions of scopolamine impairs spatial discrimination learning in rats (Whishaw, 1985; Blokland et al., 1992), and in humans, scopolamine disrupts the normal dynamics of theta oscillations in the hippocampus crucial for normal learning and memory (Gedankien et al., 2023). However, scopolamine is not selective for postsynaptic receptors; it could have also blocked muscarinic receptors located presynaptically on cholinergic terminals. These are known to act as inhibitory autoreceptors which increase acetylcholine release (Raiteri et al., 1989; Scali et al., 1995; NordstrÖM and Bartfai, 2008). There is some evidence that higher acetylcholine release, especially in the prefrontal cortex, improves attentional performance in rats (Himmelheber et al., 2000; Passetti et al., 2000; Dalley et al., 2001). Enhanced attentional processing could, ostensibly, explain why males in the current study, regardless of age, outperformed the females in discrimination learning following chronic injections. Although male monkeys show faster and more accurate learning than females on certain cognitive tasks (Lacreuse et al., 2005; Workman et al., 2019; Rothwell et al., 2022), this advantage declines with age (Lacreuse et al., 1999). Male marmosets are known to be easily distracted and sensitive to reward omissions relative to females (LaClair and Lacreuse, 2016). An attentional boost with acetylcholine could feasibly benefit the males in their performance.

The cognitive-behavioral effects of scopolamine have been well studied in rodents, monkeys and humans (Miravalles et al., 2025), but the findings are often mixed even though the underlying pharmacology is consistent. Our data suggest that individual variability in age and sex has a strong impact on overall experimental results; important biological patterns can often be masked when data is summarized into averages. This is particularly important in small-sample studies, where one or two outliers can disproportionately shift the mean. Mechanistic insights through scopolamine induced neural activity, neuromodulatory interactions (e.g., with dopamine) and bridging age and sex effects will provide fundamental insights into modeling the key aspects of cholinergic system dysfunction which is implicated in several disorders of mental health including schizophrenia, depression and aging.

## Acknowledgements

This research was supported by the Intramural Research Program of the National Institute of Mental Health (ZIA MH002951 to Y.C), part of the National Institutes of Health (NIH). The contributions of the NIH author(s) are considered Works of the United States Government. The findings and conclusions presented in this paper are those of the author(s) and do not necessarily reflect the views of the NIH or the U.S. Department of Health and Human Services. We thank Ms. Erina He from the Medical Arts Branch at NIH for help with illustrations. We are indebted to the NIMH Veterinary Medicine and Resources Branch (VMRB) and Central Animal Facility (CAF) for their services in animal health, hygiene, and husbandry.

## Supplementary materials

**Figure S1.**
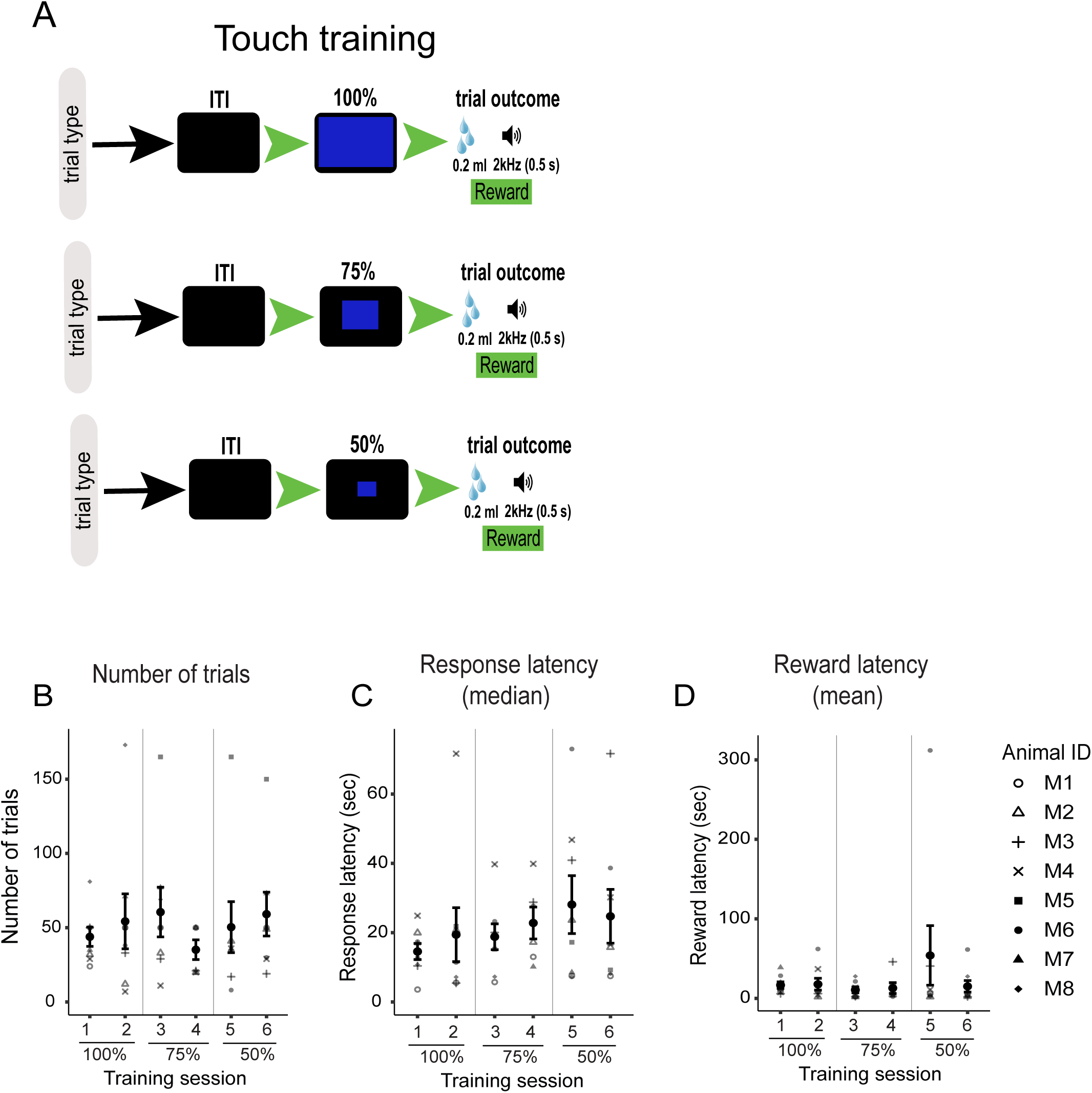
Touch training protocol and pretraining performance. **A.** Schematic illustration of how animals were trained to touch the screen. The intertrial interval (ITI) was initiated after the reward was collected but the duration of the ITI varied from trial to trial since animals returned to their homecage before initiating the next trial. After the ITI, a blue rectangle was presented which occupied 100% of the screen. A touch anywhere on the screen resulted in a 0.2ml reward and a 2kHz sound for 0.5 sec. Over successive days, the size of the blue rectangle was gradually reduced to 75% and then 50% so that it was centrally located on the screen to hone the animal’s dexterity. The touch sensitive area was the same size as the rectangle. **B.** Graph shows the number of trials committed by animals on average (± S.E.M.) across training days when the rectangle occupied 100%, 75%, and 50% of the touch sensitive space. **C.** Mean (± S.E.M.) response latency training days. Median values were used since animals voluntarily chose to enter the test box within the 2 hr training session which varied the mean from trial to trial. **D.** Graph shows reward collection latency across training days.

